# Genetic or pharmacological disruption of the MSH3 Y245/K246 IDL binding pocket slows CAG repeat expansion

**DOI:** 10.64898/2026.02.26.707948

**Authors:** Rob Goold, Jasmine Donaldson, Florence Gidney, Philip Goff, Joseph Hamilton, Marwa Elmasri, Lucy Coupland, Michael Flower, Sarah J Tabrizi

## Abstract

Recent genetic studies have shown somatic expansion of the CAG repeat is the key process driving Huntington’s disease (HD) pathogenesis. Recognition of insertion deletion loops (IDLs), lesions prone to form within the CAG repeat, by Mutsβ (MSH3/MSH2) is thought to be the primary event in the expansion process. This starts a cascade that leads to error prone repair and incorporation of additional CAG units into the repeat. *In vitro data shows* MSH3 binds IDLs through a DNA binding pocket formed by MSH3 residues Y245/K246. In this study, we investigated the significance of this DNA binding motif in CAG repeat expansion using cell lines harboring long, unstable *HTT* CAG repeats. Genetic disruption of the MSH3 Y245/K246 motif significantly reduced DNA interaction, exhibited MMR deficiency in a frameshift mutator assay and abrogated repeat expansion in a U2OS cell line expressing mutant *HTT* exon 1. Pharmacological blockade of this site using a small molecule targeting the DNA binding pocket similarly reduced DNA binding and repeat expansion in a U2OS cell line. Crucially, this molecule also slowed CAG repeat expansion in medium spiny neurons derived from HD patient-iPSCs. Targeting of the MSH3 IDL binding pocket may represent a possible therapeutic strategy.

## Introduction

Huntington’s disease (HD) is a devastating genetic neurodegenerative disorder characterized by progressive deterioration of movement, cognition, and mental health. The root cause of HD is an unstable and expanded CAG trinucleotide repeat within the *HTT* gene (1). This region typically encodes a polyglutamine (polyQ) tract of approximately 20 glutamines in the general population. However, a repeat length of 36 CAGs or more is associated with disease, with longer repeats correlating with earlier disease onset and increased severity.

The CAG repeat is known to be somatically unstable, meaning it can expand during an individual’s lifetime, leading to very long repeats (exceeding 1000 CAGs) observed in post-mortem brain tissue of HD patients (2,3). This expansion is tissue-specific, with more pronounced increases in repeat length found in brain regions primarily affected in HD, particularly the striatum (2). Interestingly, experimental HD models with CAG repeat lengths in the 40s, typically associated with human disease, often fail to exhibit a clear phenotype. This has led to the hypothesis that a significant expansion of the trinucleotide repeat, generating a longer polyQ sequence in the Huntingtin protein, is essential for disease manifestation. A proposed threshold for cellular toxicity is greater than 150 repeats (4,5). The expanded polyQ region in the mutant Huntingtin protein (mHtt) gains a toxic function, disrupts various cellular processes and ultimately leads to cell death (1). Therefore, somatic instability (SI), or the expansion of the CAG repeat, is a critical pathogenic process in HD.

Genetic studies have implicated DNA repair proteins as crucial modifiers of HD(6). Genome-wide association studies (GWAS) have identified FAN1, a DNA nuclease involved in interstrand crosslink (ICL) repair and replication fork dynamics (7,8), as a major regulator of HD age at onset (AAO) (9,10). Subsequent research has demonstrated that FAN1 slows SI in both cell and mouse models, suggesting this activity underlies its modulatory role (11,12). The GWAS analysis also identified components of the DNA mismatch repair (MMR) pathway, specifically MSH3, MLH1 PMS1 and PMS2, as regulators of HD progression (9,10). Earlier mouse studies had already pointed to MMR as a driver of both HD phenotypes and repeat expansion (13,14), providing a plausible mechanism for their role in the disease. These findings highlight that DNA damage response (DDR) proteins can either protect against or exacerbate the HD mutation through their modulation of CAG repeat length (6).

MMR is generally a protective mechanism, scanning the genome for base pair mismatches or small insertion-deletion loops that can arise during DNA replication and transcription (15). MMR is initiated by one of two heterodimeric MutS homolog complexes: MutSα (MSH2/MSH6) and MutSβ (MSH2/MSH3). MutSβ (MSH2/MSH3) favours small, extra-helical loops (insertion-deletion loops or IDLs), while MutSα (MSH2/MSH6) prefers single base mismatches. The recognition of an appropriate lesion by either MutS complex triggers the recruitment of MutL protein heterodimer complexes, forming ternary complexes. MLH1 is a common component in all MutL heterodimers, partnering with PMS2 (MutLα), PMS1 (MutLβ), or MLH3 (MutLγ) (15). Upon activation by interaction with MutS and PCNA, MutLα creates endonucleolytic incisions in the DNA flanking the mismatch (16), while MutLγ nicks the opposing strand (17). PMS1 lacks a nuclease domain, so MutLβ lacks endonuclease activity, and its specific function remains unknown. The lesion is subsequently removed by exonuclease EXO1, followed by resynthesis and ligation facilitated by DNA polymerase δ and LIG1 (15).

Less is understood about the mechanisms driving SI in the presence of a mutant CAG repeat, where MMR paradoxically adopts a detrimental role. In this context, aberrant attempts by MMR components to repair DNA lesions lead to the inclusion of additional CAG units into the repeat. While the precise mechanism remains unclear, this process is dependent on MutSβ, a dependency reflected in GWAS findings and studies in HD cell and animal models (18-23). In contrast, MutSα plays a limited role in repeat expansion (22,23). *In vitro* studies have shown that MutSβ binds with high affinity to IDLs, including those formed by short (1-3) CAG loopouts (24-26). These IDLs are prone to form in highly repetitive sequences such as the *HTT* CAG repeat, a phenomenon attributed to strand slippage, polymerase stalling, and the formation of R-loops during transcription or replication. The recognition of these lesions by MutSβ and the subsequent recruitment of downstream factors like MutL complexes and Exo1 result in the aberrant repair pathway that ultimately drives repeat expansion. Notably, all MutL factors contribute to CAG repeat expansion (14,27), and knocking down any single component (MLH1, PMS1, PMS2, or MLH3) slows expansion in an *ex vivo* HD model (28). Thus, MSH3/MutSβ, through its DNA binding and protein-protein interactions (PPI), plays a central role in SI (15).

MSH3 interacts with MSH2 to form the functional MutSβ heterodimer. This interaction is mediated by two primary regions: an N-terminal region (amino acids 126-250) and a carboxy-terminal region (amino acids 1050-1128) (29). MSH3 binds to MLH1 and PCNA via overlapping motifs located near the N-terminus. PCNA binds through a conserved PIP box spanning residues ^21^QAVLSRFF^28^, while MLH1 binds to the SRFF MIP box sequence contained within the PIP box. *In vitro* data suggest that these two proteins compete for occupancy of these sites (30). Additionally, MSH3 possesses intrinsic ATPase activity, which is essential for MMR and has been linked to repeat expansion (31).

MSH3’s binding to IDLs is mediated by a binding pocket within its mismatch binding domain (MBD). Structural studies have shown that tyrosine 245 interacts with the base pair immediately upstream of the IDL, and lysine 246 makes contact with the phosphodiester backbone of the IDL (25,32). Substitutions at these positions (Y245S/K246E) in MSH3 abolish the interaction of MutSβ with insertion/deletion loops, hairpin loops, and G4/R-loops (33,34), all structures implicated in trinucleotide repeat expansion.

In this study, we investigated the significance of the MSH3 Y245/K246 DNA binding motif in CAG repeat expansion using cell lines harboring long, unstable *HTT* CAG repeats. Genetic disruption of this motif significantly reduced DNA interaction and abrogated repeat expansion in a U2OS cell line expressing mutant *HTT* exon 1. This MSH3 mutant also exhibited MMR deficiency in a frameshift mutator assay. Furthermore, pharmacological blockade of this site using a small molecule targeting the DNA binding pocket (35) similarly reduced DNA binding and repeat expansion in a U2OS cell line. Critically, this molecule also slowed CAG repeat expansion in medium spiny neurons (MSNs) derived from HD patient-induced pluripotent stem cells (iPSCs).

## Materials and Methods

### U2OS Cell culture and manipulation

U2OS cells featuring FRT sites introduced into the genome were a gift from Prof. John Rouse (University of Dundee, Scotland). Routine culture of this line, transduction with a lentiviral *HTT* exon 1 construct harboring 118 CAG repeats (LV HTT exon 1 118Q) and expansion assays were previously described (12). *MSH3* KO and introduction of transgenes into the FlpIn site were performed as previously described (12,21). Myc MSH3 6a cloned into the pcDNA 5.1 FRT/TO.puro vector was generated by VectorBuilder. Mutations were introduced into this backbone by SDM using the QuickChange XL kit (Agilent, CA, USA). Two lines of U2OS MSH3^-/-^ cells were isolated and complemented with the myc MSH3 forms. These were used in parallel for all the experiments described in the manuscript. For example, expansion assays were performed in quadruplex, with two independent timecourses from each KO clone or complemented derivative. Transgene expression was induced by adding doxycycline (dox) direct to cell media routinely at 0.1 ng/ml for endogenous expression levels or at 10 ng/ml for overexpression studies. When using the MSH3^E976A^ mutant 1 ng/ml doc was used to induce endogenous levels of MSH3. Fresh media and dox were added every two or three days. CP1 (2-chloro-N-[4-methyl-5-[(4-methylphenyl)methyl]-1,3-thiazol-2-yl]acetamide) was purchased from ENAMINE and added to cell media from a 20 mM stock dissolved in DMSO. Media with fresh CP1 was replaced every two days. Toxicity of CP1 was assessed using an MTT viability assay following CP1 exposure for 24 or 48 h as previously described (12,21).

### Human iPSC line maintenance and differentiation

The QS5.1 125 CAG iPSC line (IRB number: CENSOi019-A) was derived previously from fibroblasts from a 7-year-old female pediatric HD individual with fully informed parental consent and ethical approval from the local ethics committee with good clinical practice (GCP) compliance. The fibroblasts were reprogrammed by Sendai-based methods at Censo Biotechnologies. This iPSC line originally had a repeat sequence containing 125 uninterrupted *HTT* CAG repeats, followed by a single CAACAG cassette. Prior to differentiation, HD iPSCs were maintained in Stem Flex (Gibco) on Geltrex (1:100, Gibco)-coated plasticware and passaged at 70-80% confluency using ReLeSR (STEMCELL technologies).

Differentiation into striatal neurons was achieved after the 36-day protocol as described in Arber *et al* (36). which involves the promotion of lateral ganglionic eminence fate via activin A treatment, followed by terminal differentiation into striatal neurons. Cultures were treated continuously with CP1 from day 36 of the differentiation process and wells were harvested every 3 to 4 weeks to monitor CAG repeat instability. Toxicity of CP1 was assessed using an MTT (Sigma-Alrich, <u>475989</u>) viability assay following CP1 exposure for 12 weeks. For immunocytochemistry cells were fixed with 4% PFA for 15 mins, washed with PBS and permeabilised with ethanol (15 mins at room temperature, RT). The cells were blocked for 1 hour in 5% BSA, 1% goat serum, 0.5% Triton X-100, (block buffer, BB), then primary antibody was added overnight in BB. After 4 washes of 10 min with PBS, samples were incubated with secondary antibodies in BB and Hoechst counterstain for 2 h RT. Following 4 further 10-min PBS washes, samples were prepared for microscopy with 90% glycerol and 0.5% N-propyl gallate in 0.1 M Tris pH 7.4. Images acquired on an Opera Phenix (PerkinElmer/Revvity) from 10 random fields in 3 wells from each experiment. Z-stacks of 10 layers were combined for each image. Primary antibodies used were NeuN (ab104224), beta III-Tubulin (ab107216), DARPP32 (ab40802), FOXP1 (ab32010) from Abcam and MAP2 (NB300-213, Novus). AlexaFlour secondary antibodies were from Invitrogen

### Immunoprecipitation, ChIP and biotinylated oligo pull down

Extracts for immunoprecipitation (IP) were prepared as previously described (12,21) and IP was done overnight at 4°C using anti-MSH3 or anti-MLH1 antibodies (BD Biosciences) coupled to protein A/G magnetic beads or anti-myc tag Dynabeads (Pierce). Biotinylated oligoes were purchased from IDT and annealed by heating to 95°C for 5 minutes then slow cooling to room temperature. Oligo sequences were:

CGACTTCCGGTAGCACGTAGCACTGCTGCGCCACGAACTGCACTCTAGGC GCCTAGAGTGCAGTTCGTGGCGCAGCAGCAGCAGTGCTACGTGCTACCGGAAGTCG

These were coupled to MyOne Streptavidin T1 Dynabeads via a 5’ biotin moiety conjugated to the top strand according to the manufacturers instructions (Pierce) using 75 ng of annealed oligo per µl of beads. Cell extracts were prepared by lysing cells in TBS pH 7.4 with 1% Triton X100 and protease inhibitors (Sigma). Lysates were sonicated for 30 seconds at 40% power in a Ultrasonic processor 500W probe sonicator then centrifuged at 20,000g for 3 min. Supernatants were adjusted to 2 mg/ml and used as inputs for pull downs, using 800 µg protein and 20 µl beads for each. Input and IP or pull down fractions were prepared for immunoblotting as previously described (12,21). Antibodies used were: MSH3 rabbit polyclonal (Proteintech, used for ChIP and some Westerns), MSH3 mouse monoclonal, MLH1 mouse monoclonal and MSH6 mouse monoclonal (BD Biosciences), MSH2 and PCNA rabbit (Cell Signaling Technology), and an actin mouse monoclonal (Sigma). ChIP analysis was performed with the EZ-Magna ChIP Kit as previously described (12,21). Results were expressed as fold increases relative to U2OS MSH3^−/−^ ChIP or control samples as indicated in the main text.

### Somatic instability

Expansion assays and fragment analysis were performed as previously described (12,21) using GeneMapper v6. software (Thermo) to align the chromatographs. To calculate modal CAG repeat length, GeneMapper data was exported and analyzed with a custom R script, available at https://caginstability.ml.

### Long-read repeat sizing

Expansion of the CAG repeat in striatal cultures was assayed using long-read repeat sizing. Genomic DNA (25 ng per reaction) was amplified using locus-specific primers flanking the HTT exon 1 CAG repeat (MF20230707_20 assay). Each primer incorporates a unique 16-bp barcode, enabling asymmetric dual-barcode sample indexing from the first amplification cycle. Primers were designed in SNP-free, haplotype-conserved regions to ensure universal amplification without allele-specific bias. The ∼3 kb amplicon minimises preferential amplification of shorter alleles by reducing the relative length difference between expanded and non-expanded alleles to approximately 5%.

PCR was performed in 20 µL reactions using Platinum SuperFi II DNA Polymerase Master Mix (Invitrogen, 12368010) with 0.3 µM of each barcoded primer under the following conditions: 98 °C for 30 s; 28 cycles of 98 °C for 10 s, 60 °C for 10 s, and 72 °C for 2 min; followed by 72 °C for 5 min. Products were verified by 1% agarose gel electrophoresis, quantified by Qubit dsDNA HS assay, normalised to equimolar concentrations, and pooled. Pooled amplicons were purified using 1.8x AMPure XP beads (Beckman Coulter, A63881).

SMRTbell libraries were prepared from pooled amplicons using the SMRTbell Prep Kit 3.0 (PacBio, 102-141-700). Amplicons underwent DNA damage repair and end-repair/A-tailing, followed by ligation of indexed SMRTbell adapters. Libraries were treated with Exonuclease III and VII to remove unligated fragments, purified with SMRTbell cleanup beads, and sequenced on the PacBio Revio platform to generate high-fidelity (HiFi) circular consensus reads.

### Bioinformatic analysis

HiFi reads were demultiplexed by dual-barcode assignment using Ophelia (https://github.com/mike-flower/ophelia) and analysed using the Duke pipeline (https://github.com/mike-flower/duke. Reads were aligned to the amplicon reference sequence and CAG repeat tract length was measured for each read. Reads were assigned to alleles based on repeat length, and the modal repeat length was determined as the most frequently observed value within predefined ranges (normal: 0-35 CAG; expanded: ≥36 CAG).

Somatic instability metrics were calculated relative to the modal repeat length of the baseline timepoint. The instability index was defined as the mean absolute deviation from the modal length; the expansion index as the mean deviation for reads above the modal length; and the expansion ratio as reads above the modal length divided by reads at the modal length. For analyses of MSN-enriched cultures presented in Figure 5, the instability index was used as the primary quantitative summary of somatic CAG repeat expansion, as it captures shifts across the full repeat-length distribution over time.

### EMAST reporter assays

Cells were transduced with a lentiviral EMAST frameshift reporter construct (37) and cultured for five days, including 0.1 ng/ml dox in complemented cell cultures. NanoLuc and Luc2 signals were measured using the NanoGlo kit (Promega) and luciferase assay substrate (Abcam) respectively. Results are expressed as NanoLuc/Luc2 normalised to WT cells

### Quantification and statistical analysis

CAG expansion time courses were analyzed by linear regression and slopes statistically compared by one-way ANOVA. Multiple comparisons were corrected for with a False Discovery Rate (FDR) of 5%. The Brown-Forsythe test was routinely used to check for homogeneity of variance. All statistical information can be found within figure legends.

## Results

### Generating a cell system to study MSH3 function in CAG repeat expansion

We have previously used U2OS cells transduced with an HTT exon1 118Q construct to study repeat dynamics and shown these cells support expansion in culture, adding approximately one CAG to the repeat every 18-20 days (12). Following MSH3 KO these cells did not support CAG repeat expansion in line with findings from other HD models (21). These cells were complemented with a WT myc tagged MSH3 6a construct expressed under a doxycycline (dox) inducible promoter (The MSH3 6a isoform is the most common allele - Ref sequence <u>NC 000005.10</u> (38)). Maintaining myc MSH3 6a construct expression at similar levels to endogenous MSH3 with 0.1 ng/ml dox rescued repeat expansion to rates seen in U2OS WT cells (Figure 1A and B). Increasing dox to 10 ng/ml lead to 8 fold overexpression of myc MSH3 6a and increased the expansion rate of the CAG repeat in the HTT exon1 118Q construct to above that seen in U2OS WT cells, approaching the levels seen in U2OS FAN1 KO cells (12) (Figure S1) . These data demonstrate the importance of MSH3 in driving CAG repeat expansion and show the U2OS system can provide a test bed to assess the role of MSH3 functions in CAG repeat expansion.

**Figure 1.**
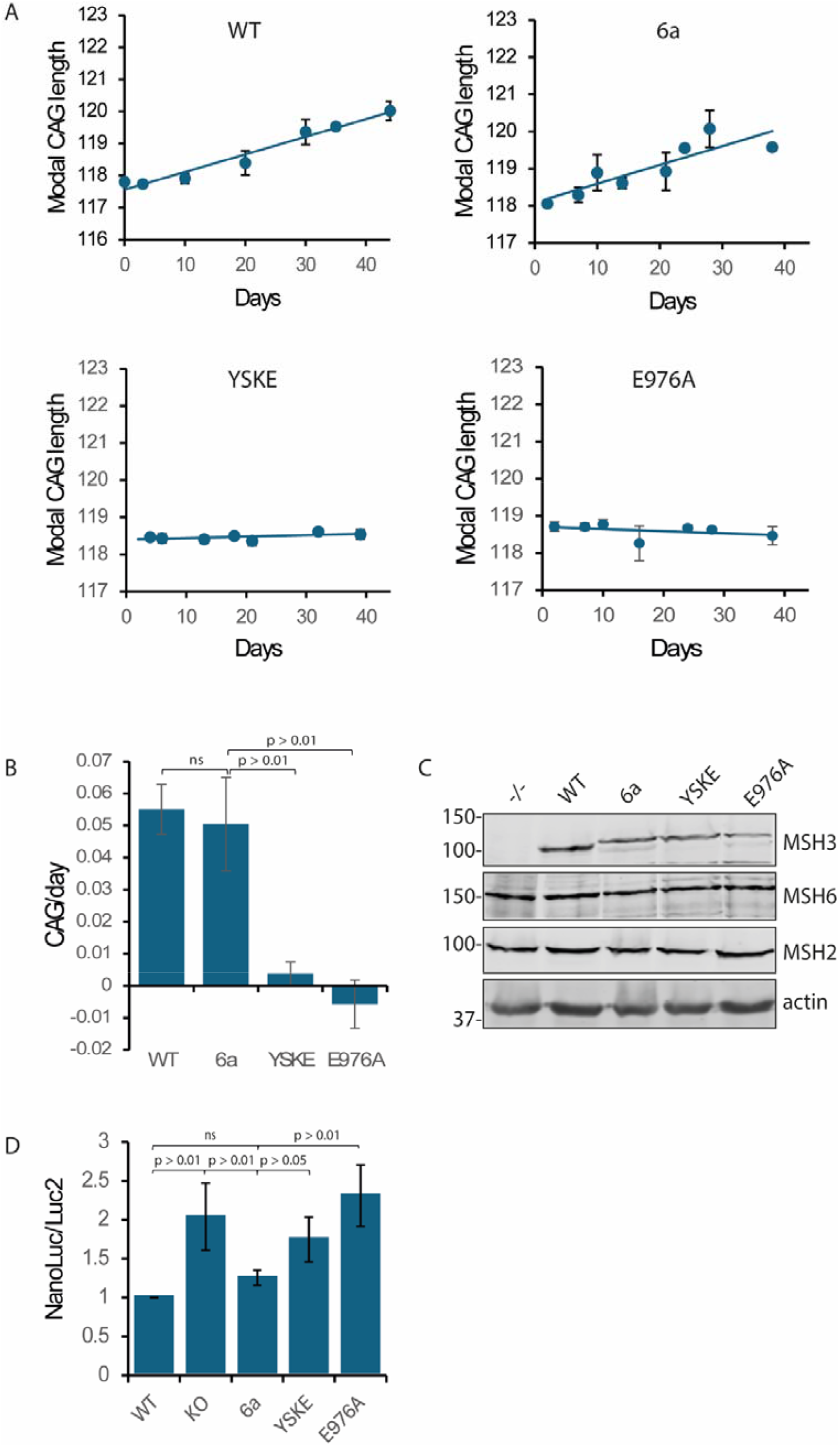
Genetic blockade of the MSH3 IDL binding pocket disrupts MMR function. **(A)** WT or MSH3 KO cells complemented with myc MSH3 6a, Y254S/K255E or E976A constructs were transduced with LV *HTT* exon 1 118 CAG. Repeat size was monitored over 40 days in culture using fragment analysis. WT cells show a smooth increase in CAG repeat size over time in culture. MSH3 KO cells do not support CAG repeat expansion (not shown). CAG repeat expansion is rescued by expression of the MSH3 6a construct (the full-length WT form) expressed at similar levels to the endogenous protein (with 0.1 ng/ml dox treatment). In contrast, cells expressing myc MSH3 Y254S/K255E or E976A mutants do not support CAG repeat expansion. **(B)** Histogram shows regression analysis of expansion curves in (A), rate CAG per day (mean ± standard error n=4, p values one-way ANNOVA). **(C)** Western blot showing expression of MutS proteins in MSH3 KO, WT and KO cells complemented with myc MSH3 6a, Y254S/K255E or E976A constructs. Complemented cells were treated with 0.1 ng/ml dox except the E976A line in which 1 ng/ml dox was used. Note the myc tagged MSH3 forms run at a higher molecular weight. **(D)** U2OS WT, MSH3 KO or MSH3 KO cells complemented myc MSH3 6a, Y254S/K255E or E976A were transduced with an EMAST frameshift reporter construct. This 16 tetranucleotide repeat construct measures MSH3 dependent MSI in cells, assayed by measuring NanoLuc signal normalised to a constitutive Luc signal. Results are expressed as NanoLuc/Luc2 normalised to WT cells (n=4 or 5 ± sd, p values one-way ANNOVA).

To test if IDL binding by MSH3 DNA binding motif plays a role in CAG repeat expansion we introduced Y254S/K255E substitutions into the 6a construct (corresponding to Y245S/K246E in the canonical human MSH3 sequence described by Gupta et al., 2012). We also generated a MSH3 E976A Walker B ATPase mutant as a comparator. This mutation is well characterised, is known to disrupt MMR activity and has been associated with reduced repeat expansion (31). Unlike the MSH3 6a construct, these mutants did not promote repeat expansion, despite expression at comparable levels (Figure 1A, B and C).

As an independent measure of MMR activity in cells expressing these mycMSH3 constructs we performed a frameshift mutator assay designed to measure MSH3 activity. This assay is based on a 16 repeat tetranucleotide sequence that is prone to Elevated microsatellite alterations at selected tetranucleotide repeats (EMAST) linked to a reporter in an expression cassette. When introduced into a cell it is prone to alterations in size, either by addition or deletion of an AAAG four nucleotide unit. This shifts the construct into frame with the reporter, in this case a NanoLuc sequence. Maintenance of the repeat region relies largely on MSH3 activity and deficiency in function increases EMAST which leads to a reporter expression (37). This EMAST reporter is preferentially sensitive to MutSβ (MSH2/MSH3) function and is less dependent on MutSα (MSH2/MSH6), making it well suited to assess MSH3-specific mismatch repair activity in this system.

In U2OS WT cells a baseline NanoLuc signal is detected after five days, indicating a relatively low level of frameshifts in the reporter construct as would be expected (37). MSH3 KO cells show a consistent increase in NanoLuc activity, roughly doubling the signal over the course of five days in culture. Importantly, complementation of the MSH3 KO cells with myc MSH3 6a, expressed at endogenous levels restores baseline EMAST activity (Figure1D).This demonstrates the MSH3 dependence of the EMAST reporter. In contrast, cells expressing the myc MSH3^E976A^ ATPase dead form show high levels of EMAST, with similar NanoLuc signal to MSH3 KO cells. Cells expressing myc MSH3^Y254S/K255E^ mutant also show EMAST deficiency but the NanoLuc signal does not reach KO levels (Figure1D).

### MSH3 molecular interactions underlie cell phenotypes

These assays show interesting MSH3 related cell phenotypes. To understand the mechanism underlying these phenotypes we have studied the molecular interactions of WT MSH3 and the myc tagged 6a and Y254S/K255E constructs.

MMR complex formation was studied using an IP approach. Using 0.1 ng/ml dox induction and MSH3 antibodies allowed a comparison of the activity of WT MSH3 and myc MSH3 6a or Y254S/K255E transgenic forms at equivalent expression levels (Figure 2 A). The procedure immunoprecipitated robust levels of MSH3 from all the input fractions. MSH2 co-fractionates with MSH3 in all the IP samples indicating the MutSβ complex formed normally in with transgenic MSH3 proteins (Figure 2 B). MLH1 cannot be detected in the IP fractions under these conditions, possibly because of the relatively low levels of MSH3 and the transient nature of the MutS/MutL interaction. The specificity of the procedure is demonstrated by use of an MSH3 KO line as control. A reverse IP using MLH1 antibodies pulls down high levels of MLH1 from all IP samples (Figure S1D). Low levels of MSH2 and MSH3 can be detected in the WT and myc MSH3 6a MLH1 IP fractions. MSH3^Y254S/K255E^ levels in MLH1 IP fractions were lower. Although this pattern of IP co-fractionation was reproducible quantifying differences in binding activity between MLH1 and the MutS components in these samples was difficult because of the low signals.

**Figure 2.**
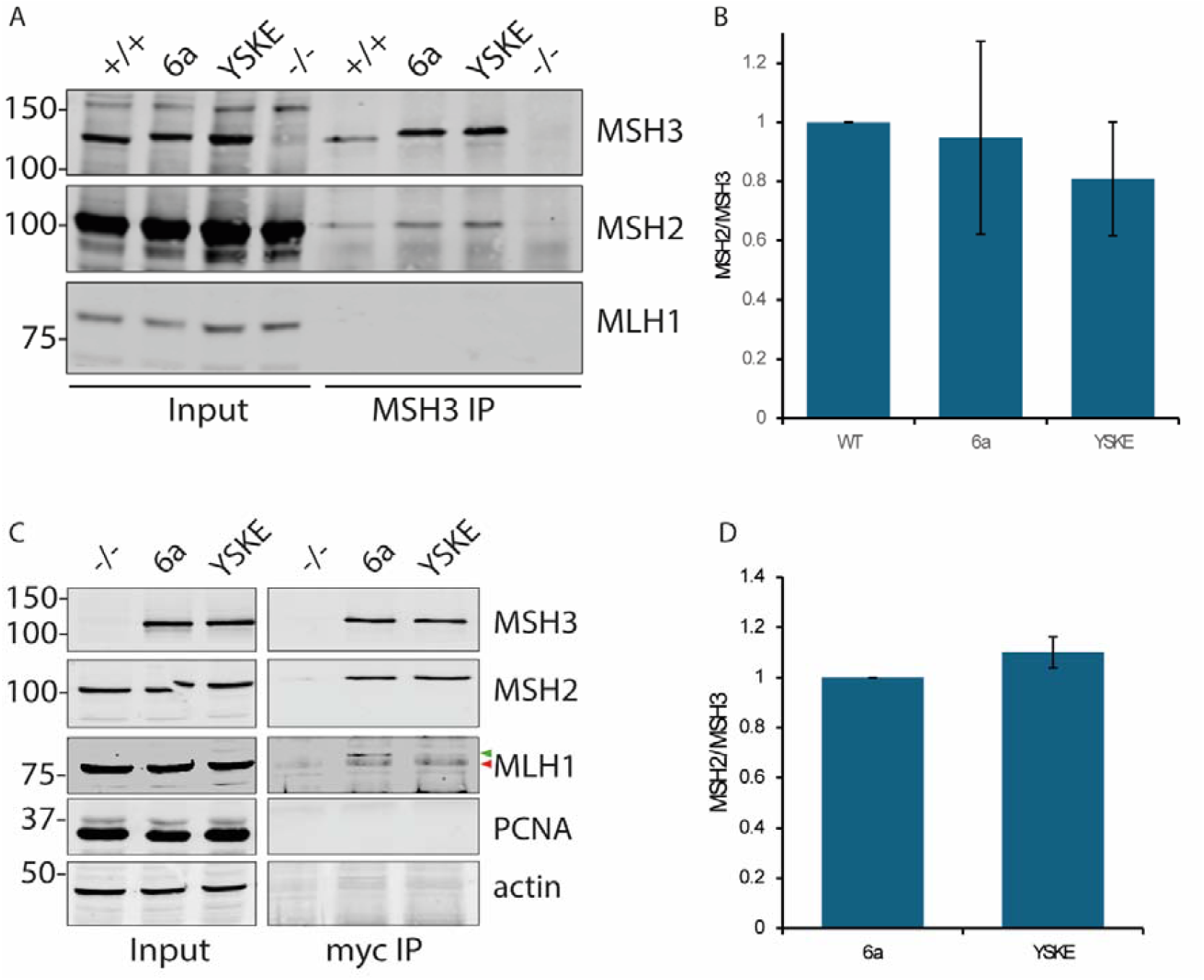
MSH3 protein interactions in WT and myc MSH3 complemented cells. **(A)** U2OS cell extracts from WT (+/+) MSH3 KO (-/-) or KO cells complemented with myc MSH3 6a or myc MSH3 Y254S/K255E (YSKE) constructs treated with 0.1 ng/ml dox were incubated O/N with MSH3 antibodies and protein A/G magnetic beads. Beads were isolated and washed with a magnetic device. Input (5%) proteins eluted from the beads were immunoblotted with the indicated antibodies. Similar levels of MSH2 were recovered in the MSH3 IP fraction from both WT MSH3, myc MSH3 6a and YSKE samples. MLH1 was not detected in the IP fractions under these conditions. **(B)** Quantification of MSH2/MSH3 binding from IP experiments normalized to WT MSH3 is shown in the histogram (n=3 ± sd). **(C)** U2OS cell extracts from MSH3 KO or KO cells complemented with myc MSH3 6a or myc MSH3^Y254S/K255E^ were incubated O/N with myc magnetic beads. Transgenes were over-expressed by approximately 8-fold with with 10 ng/ml dox. Beads were isolated and washed with a magnetic device. Input (5%) and proteins eluted from the beads were immunoblotted with the indicated antibodies. Similar levels of MSH2 were recovered in the myc IP fraction from both MSH3 6a and YSKE samples indicating the MSH3 YSKE mutant binds MSH2 normally. Low levels of MLH1 (green arrowhead) were consistently detected in myc MSH3 6a IP fractions. Note a non-specific band running slightly faster than MLH1 (red arrowhead) is detected in all of the IP fractions **(D)** Quantification normalized to MSH3 6a is shown in the histogram (n=3 ±sd).

One advantage of the U2OS system is that the level of transgene expression can be precisely controlled. As we showed previously myc MSH3 6a overexpression accelerates CAG repeat expansion (Figure S1). Expansion is generally believed to be dependent on MutSβ driven MMR which in turn requires ternary complex formation between DNA, MutSβ and MutL components (15). To see if MSH3 overexpression is accompanied by increased MutSβ heterodimerization and MutL binding we performed MLH1 IPs (Figure S1). In these experiments cell extracts were prepared for IP from myc MSH3 6a cells treated with 0.1 or 10 ng/ml dox. IP’s were performed with MLH1 specific antibodies and input or IP fractions were immunoblotted for MSH3, MSH2 and MLH1 (Figure S1E). Increased MSH3 expression was accompanied by a small increase in MSH2 expression but a bigger increase in MSH3/MSH2 co-IP with MLH1, suggesting an increase in MutSβ/MutL complex formation (Figure S1). MSH3 levels in the cell extracts and IP fraction increased proportionally more than MSH2, reflecting the incorporation of most of the MSH2 into MutSα which is the dominant MutS form present in cells.

This suggests that overexpression of MSH3 results in an increase in active MMR complex capable of enhancing CAG repeat expansion that retains the molecular interactions that drive the process. We took advantage of this to look at myc MSH3 protein and DNA interactions in more detail by overexpressing the transgenes in our cells; this facilitated detection of binding partners.

Firstly, we looked at myc MSH3 protein interactions using myc beads to pull down the tagged proteins. The IP fractions from the MSH3 6a and MSH3^Y254S/K255E^ forms contained equivalent levels of MSH2 showing the Y254S/K255E substitutions did not affect MutSβ heterodimerisation (Figure 2C and D). MSH3 6a IP fractions contained low but reproducibly detectable amounts of MLH1 suggesting some MutL interactions are retained. This probably reflects the transient nature of MutSβ/MutL interaction. IP fractions from MSH3^Y254S/K255E^ cell extracts contained less MLH1, though this was difficult to quantify accurately given the low signal. MutL recruitment to MutSβ takes place on a DNA substrate in situ so reduced DNA association of the mutant form may impair its ability to bind MLH1. PCNA and FAN1 were not detected in the IP fractions.

DNA interactions were initially assessed by ChIP using a specific antibody to isolate MSH3 and associated DNA. Extracts from MSH3 KO and myc MSH3 cells were prepared that gave 8-fold overexpression of myc MSH3 forms (Figure S1). This allowed unequivocal detection of CAG repeat and control HTT GFP DNA in the ChIP fractions (Figure 3). ChIP fractions from myc MSH3 6a contained more CAG repeat and HTT GFP DNA compared to background levels detected in ChIP fractions prepared from KO cells or those isolated with a non-specific IgG control antibody. In contrast, the myc MSH3^Y254S/K255E^ mutant showed reduced DNA association, either with the CAG repeat or downstream HTT GFP DNA (Figure 3 A). Quantification of the HTT GFP amplicon, chosen as it gave a clearer signal (Figure 3 B), showed significant differences in the levels of DNA in the mycMSH3 6a ChIP fractions compared to KO and myc MSH3^Y254S/K255E^. Control ChIP fractions made using RNA polymerase II antibodies contain similar amounts of DNA in all samples.

**Figure 3.**
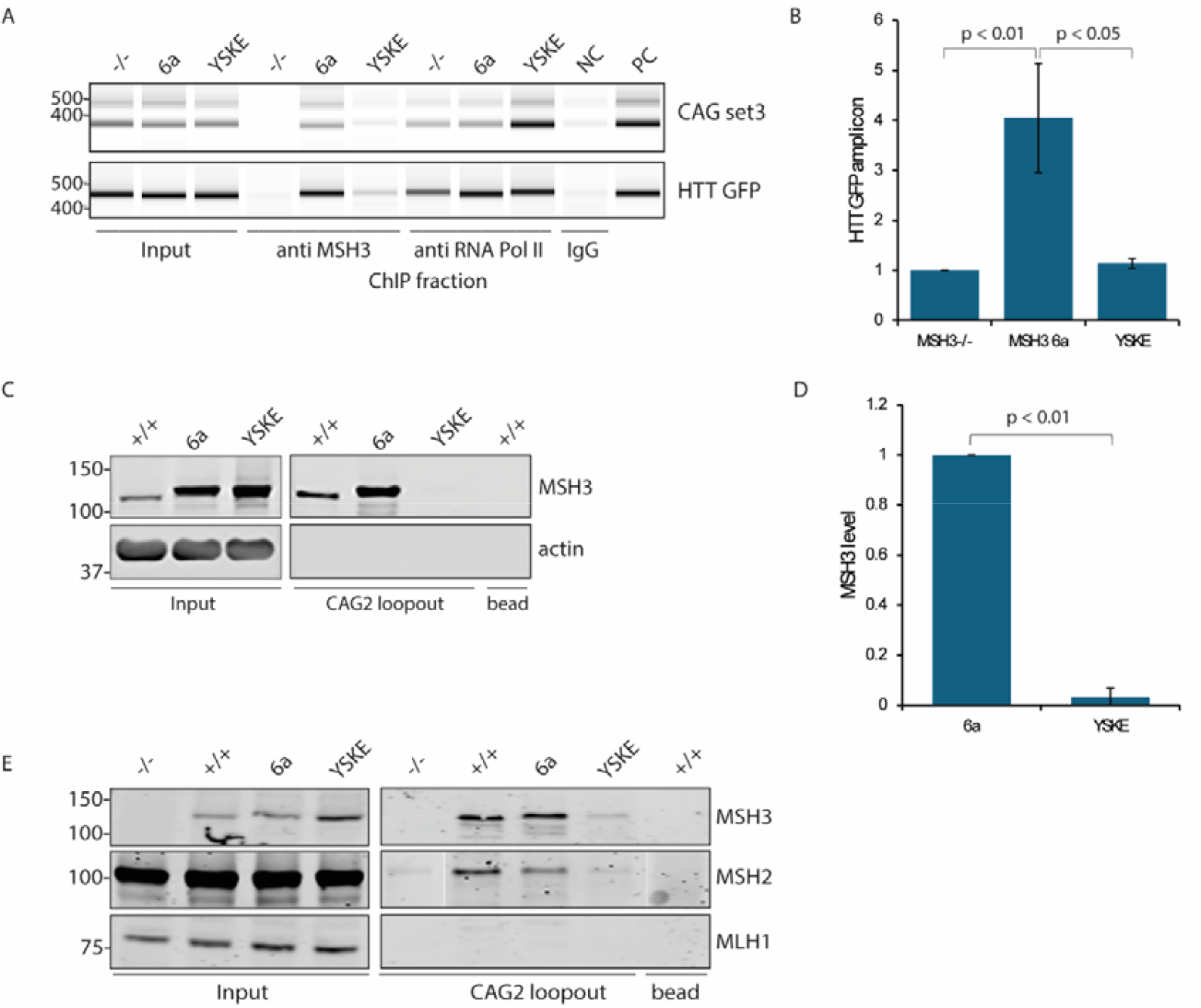
MSH3 DNA interactions in WT and myc MSH3 complemented cells. **(A)** Cell extracts from HTT exon 1 118Q transduced MSH3 KO cells or (-/-) or KO cells complemented with myc MSH3 6a or myc MSH3 Y254S/K255E (YSKE) were prepared for ChIP analysis and immunoprecipitated with anti MSH3, RNA PolII or control non-specific mouse IgG antibodies (NC). DNA from input and ChIP fractions was purified and probed with primers targeting the *HTT* exon 118Q CAG repeat in the construct (CAG set 3) or GFP primers downstream from this in the insertion cassette (HTT GFP). Input (5%) and ChIP fractions were analysed using TapeStation apparatus and software. MSH3 ChIP fractions from YSKE mutant contain less CAG repeat and HTT GFP DNA. Note the myc MSH3 cells were treated with 10 ng/ml dox which induces ∼8x MSH3 overexpression relative to WT cells. Plasmid DNA was amplified alongside the ChIP fractions as a positive control (PC) **(B)** Quantification of HTT GFP levels in ChIP fractions normalized to MSH3^-/-^ is shown (n=4 ± sd, p values one-way ANNOVA). **(C)** U2OS cell extracts from WT cells or KO cells complemented with myc MSH3 6a or myc MSH3 Y254S/K255E (YSKE) were prepared and incubated O/N with 2 CAG loopout oligos immobilized on streptavidin magnetic beads. Beads were isolated and washed with a magnetic device. Input (5%) and eluted proteins (CAG2 loopout) were immunoblotted with the indicated antibodies. Unconjugated beads (bead) were used as controls (WT sample only). Note the myc MSH3 cells were treated with 10 ng/ml dox which induces ∼8x MSH3 overexpression relative to WT cells (compare lanes +/+ and 6a). **(D)** Quantification of MSH3 levels in CAG2 loopout pull down fractions normalized to myc MSH3 6a is shown (n=4 ± sd, p values one-way ANNOVA). **(E)** U2OS cell extracts from WT cells, KO cells or KO cells complemented with myc MSH3 6a or myc MSH3 Y254S/K255E (YSKE) and treated with 0.1 ng/ml dox were prepared. These were incubated O/N with magnetic streptavidin beads conjugated to 2 CAG loopout oligos or empty beads. Input (5%) and eluted proteins (CAG2 loopout) were immunoblotted with the indicated antibodies. Equivalent levels of MSH3 and MSH2 were recovered in the pulldown fractions from WT and myc MSH3 6a indicating the WT and myc tagged MSH3 forms have similar activity. Proportionally less of the MSH2 input was pulled down compared to MSH3. Pull downs from myc MSH3 Y254S/K255E extracts contained only low levels of MSH3 and MSH2. MSH3 KO cell extracts (-/-) and unconjugated beads (bead, WT sample only) were used as controls. N.B. Lanes (-/-) and (+/+, bead) have been switched around in the middle (MSH2) panel to match the arrangement of the top (MSH3) panel.

To confirm this data we also performed affinity pull down of MSH3 from cell extracts using immobilized biotinylated DNA oligos as bait. We selected a 50-mer sequence with a two CAG loopout inserted in the 5’-3’ strand, mimicking the likely DNA lesion targeted by DNA repair apparatus during SI. WT MSH3 and myc MSH3 6a were detected in the pull downs, binding with apparent high affinity to the 2 CAG loopout oligo (Figure 3 C). Only low to background levels of myc MSH3^Y254S/K255E^ were detected in pull downs fractions, showing the mutation prevents effective interaction with the DNA oligos (Figure 3 C and D). No MSH3 was detected in empty bead control fractions. Similar results were obtained when pull downs were performed from cells expressing endogenous levels of MSH3 (Figure 3 E). MSH2 was also detected in the pull downs indicating MutSβ is binding to the oligo. Proportionally less of the MSH2 input was pulled down compared to MSH3. This is likely because most of the MSH2 is associated with MSH6 which doesn’t bind to the oligo under the conditions used for these experiments. Similarly, MLH1 was not detected in pull down fractions (Figure 3 E).

### Pharmacological blockade of the MSH3 IDL binding pocket reduces DNA interaction

To confirm the importance of the MSH3 IDL binding site in CAG repeat expansion and test its potential as a therapeutic target we have used a recently identified small molecule that binds close to Y254/K255 and blockades DNA binding (35). This molecule, 2-chloro-N-[4-methyl-5-[(4-methylphenyl)methyl]-1,3-thiazol-2-yl]acetamide (referred to as CP1 hereafter), was one of a library of cysteine targeting molecules tested. Consistent with its effect on MSH3 activity, CP1 was shown to bind irreversibly to cysteine 252, whose side chain protrudes into the DNA-binding pocket of MSH3.

To verify target engagement in our system we added CP1 to WT U2OS cell extracts for one hour and assessed MSH3 binding to a CAG loopout oligo. This procedure clearly showed dose dependent inhibition of MSH3 and MSH2 oligo binding (Figure 4 A and B). Next, we added CP1 to live cells and assessed target engagement in situ. Even at the moderate CP1 concentrations (10 µM and 20 µM) added to the lysates cell toxicity was observed. However, viability assays showed lower CP1 concentrations were well tolerated by the cells (Figure S2 A). Therefore, CP1 was added to the live cells at 2.5 and 5 µM. Cells were treated overnight then extracts were prepared for CAG loopout oligo pull down. Immunoblotting shows robust MSH3 and MSH2 binding to the oligo following vehicle (DMSO) addition. Again, MSH2 levels relative to the input fractions were proportionally less compared to MSH3. Importantly, clear dose dependent inhibition of MSH3/MSH2 binding was observed at 2.5 and 5 µM CP1 (Figure 4). Given the reduced concentration of CP1 used, inhibition was more effective after cell addition than adding CP1 direct to lysates, possibly because of the longer exposure time (Figure 4 A and B).

**Figure 4.**
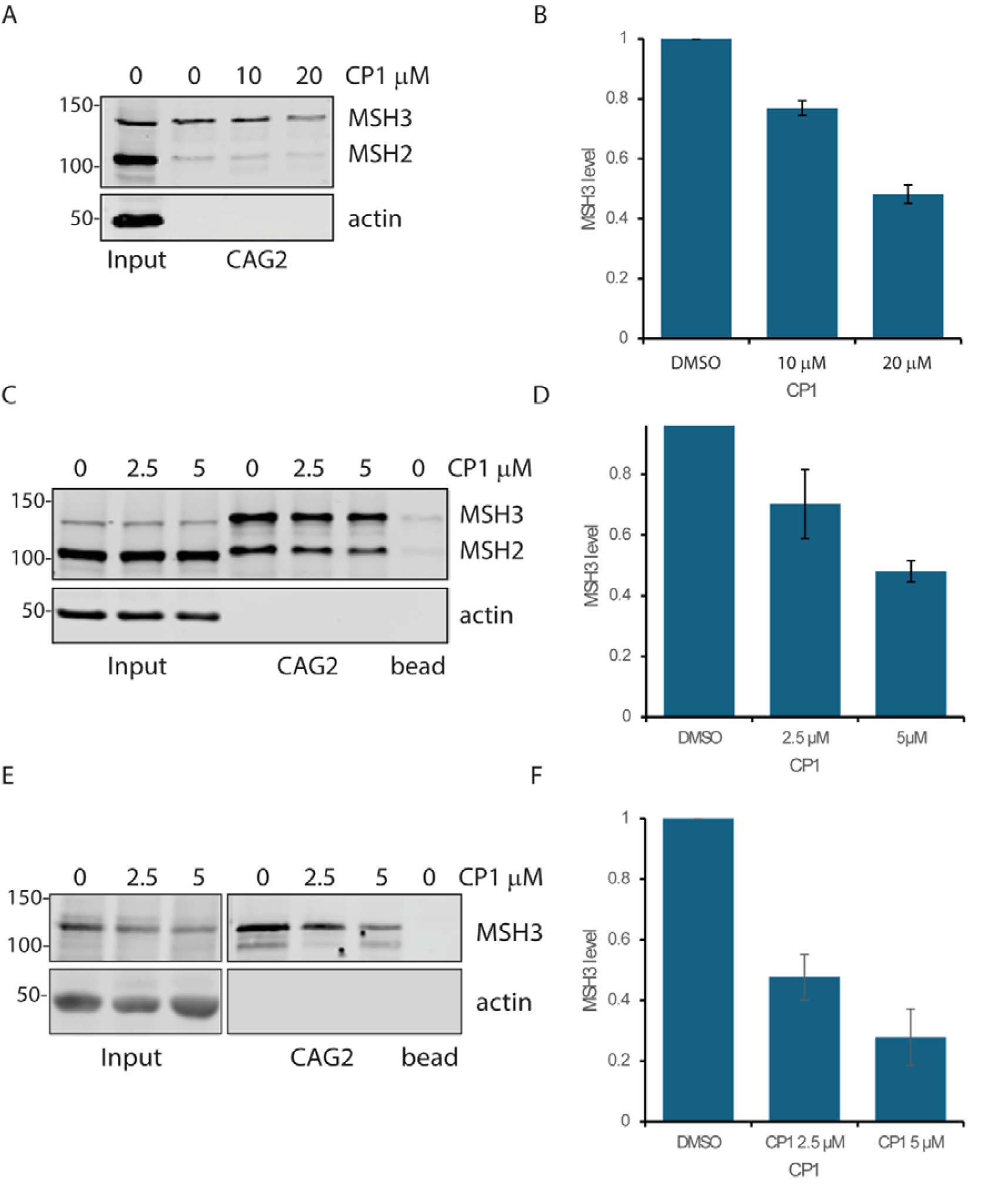
Pharmacological blockade of the MSH3 IDL binding pocket reduces DNA interactions. **(A)** U2OS cell extracts from WT cells were treated with the indicated CP1 concentrations for 1 h prior to incubation with 2 CAG loopout oligos immobilized on streptavidin magnetic beads. Beads were isolated and washed with a magnetic device. Input (5%) and eluted proteins (CAG2) were immunoblotted with the indicated antibodies. MSH3 and MSH2 were recovered in the pulldown fractions in vehicle (DMSO) treated extracts. CP1 treatment reduced the recovery. **(B)** Quantification of MSH3 levels in CAG2 loopout pull downs normalized to DMSO control is shown in the (n=4 ± sd). **(C)** U2OS WT cells were treated for 24 h with CP1 at 0, 2.5 or 5 uM and cell extracts were prepared and incubated with 2 CAG loopout oligos immobilized on streptavidin magnetic beads as described in panel (A). MSH3 and MSH2 recovery in the bead fraction was reduced by CP1 treatment. Unconjugated beads were included as a control (E). (**D**) Quantification MSH3 levels in CAG2 loopout pull downs normalized to vehicle treated cells is shown (n=4 ± sd). (**E**) HD MSNs were treated with CP1 (2.5 and 5 µM) for 48 h, cell extracts were prepared and 2 CAG loopout oligo binding assays performed as in panel (A). (**F**) Quantification MSH3 levels in CAG2 loopout pull downs normalised to vehicle treated cells is shown (n=3 ± sd).

ChIP experiments were also performed to test target engagement (Figure S2 D). FAN1 KO cells^12^ harboring the LV HTT exon 1 118Q insert were used. This allowed the interaction of MSH3 with the expanded CAG repeat to be assessed. These were treated with vehicle or 5 µM CP1 and prepared for ChIP. Endogenous MSH3 was immunoprecipitated using specific antibodies and DNA was probed with CAG repeat and HTT GFP primers as described earlier (Figure 3). Low levels of DNA were detected in DMSO treated ChIP fractions. This low level of DNA is typical of endogenous MSH3 ChIP in our hands and probably reflects both the relatively low levels of MSH3 in the cells and the transient nature of the MutSβ DNA association. In particular, the long allele of the 118 CAG insert was hard to detect, a consistent problem with the ChIP procedure, especially with cells harboring very long CAG repeats. This reflects the difficulty in amplifying the CAG repeat as it lengthens and the fact that the ChIP protocol requires DNA to be fragmented meaning longer DNA sequences are more likely to be broken during preparation. Although CP1 treatment consistently reduced the level of CAG repeat and downstream HTT GFP DNA detected in the ChIP fractions, quantification of the effect was difficult due to the low signal (Figure S2). By contrast, control ChIP experiments using an RNA polymerase II antibody contained robust levels of DNA which were not affected by CP1 treatment. Together this data shows CP1 is capable of blockading MSH3 DNA interactions at concentrations well tolerated by U2OS cells.

To test the applicability of such compound in a post-mitotic, disease-relevant cell type CP1 target engagement was also assessed in HD MSNs carrying an expanded *HTT* CAG repeat (>125 CAG) (39). In line with U2OS cell data, MSH3 interaction with the CAG2 loopout oligo was reduced by 2.5 and 5 µM CP1 treatment (Figure 4 G and F).

### Pharmacological blockade of the MSH3 IDL binding pocket slows CAG repeat expansion

To test the effect of pharmacological blockade of MutSβ DNA binding on CAG repeat expansion we treated U2OS FAN1 KO cells transduced with the HTT exon 1 118Q insert with CP1 at 2.5 and 5 µM. These cells had been cultured with the exon 1 construct for three months during which time the repeat had expanded to 127 - 128 CAGs, compared to freshly transduced cells which size to roughly 118 CAGs (Figure 1). These cells were chosen because of the fast expansion rate which reduces the time required in culture to see significant changes in CAG length. The cells were cultured with vehicle or CP1 for 20 days and sampled every 4 or 5 days during that time. Genomic DNA was purified from each sample and CAG repeat length was monitored using fragment analysis. DMSO treated cells expanded at one CAG repeat unit added per 9-10 days in line with previously published data (12). CP1 treatment slowed expansion at both 2.5 and 5 µM (Figure 5 A-C), showing dose dependency roughly in line with the target engagement (Figure 5D and 4D). Although significance was not reached at 2.5 µM CP1, 5 µM treatment significantly slowed expansion relative to vehicle.

**Figure 5.**
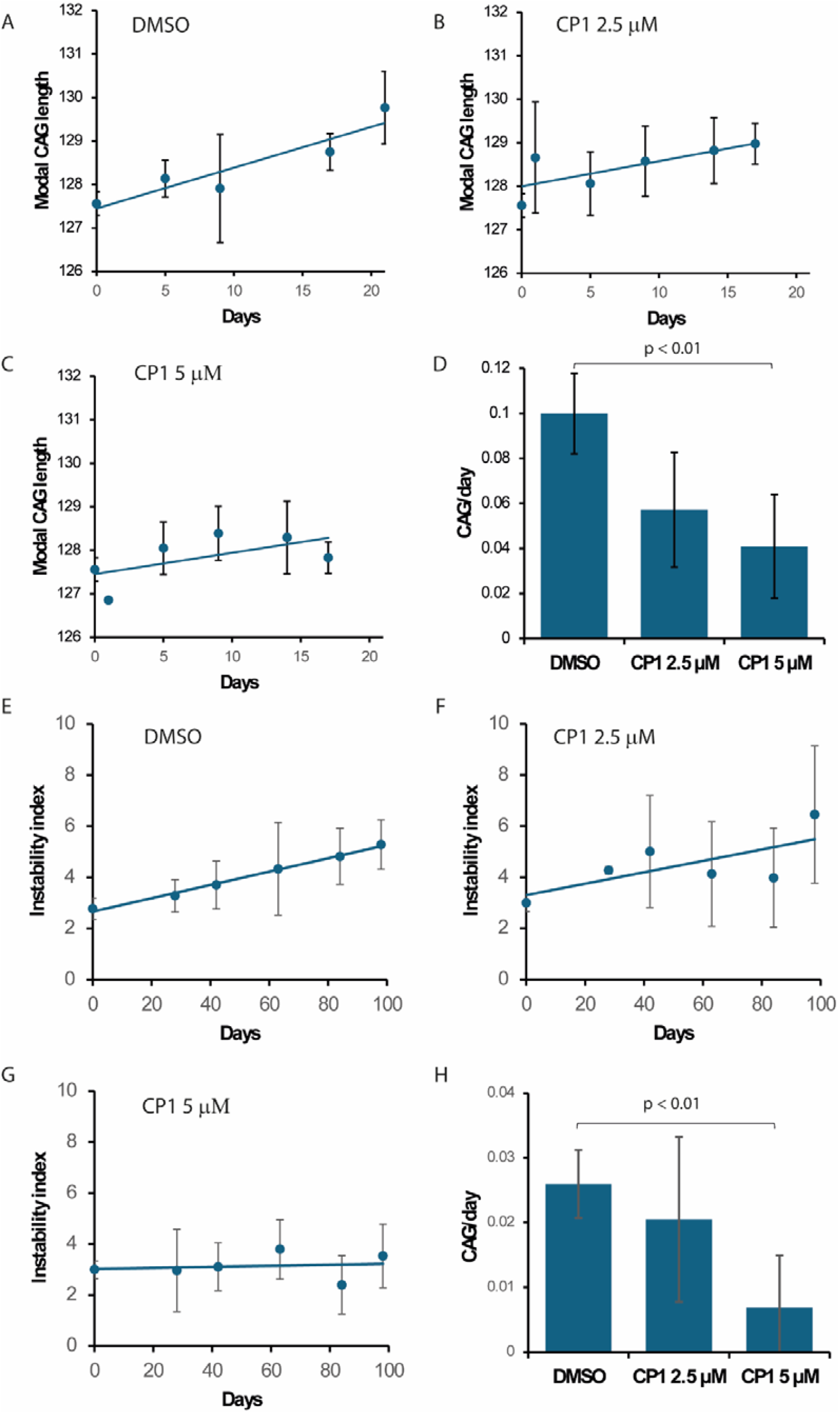
Pharmacological blockade of the MSH3 IDL binding pocket slows CAG repeat expansion. U2OS FAN1 KO cells previously transduced with LV HTT exon 1 118 CAG were cultured in the presence of vehicle (DMSO) or CP1. Repeat size was monitored over 20 days in culture using fragment analysis. Vehicle treated cells (**A**) show a smooth increase in CAG repeat size over time in culture. CP1 treatment at 2.5 (**B**) and 5 uM (**C**) progressively reduced CAG repeat expansion rate (mean ± sd, n=4). (**D**) The histogram shows regression analysis of the expansion curves shown in panels A-C (CAG per day ± standard error, n=4). Expansion rate at 5 µM CP1 is significantly slower. (**E**) Long-read repeat sizing of the endogenous HTT CAG repeat in iPSC-derived MSN-enriched cultures treated from day 36 with vehicle (DMSO). (**F**) Corresponding MSN-enriched cultures treated with CP1 (2.5 µM). (**G**) Corresponding MSN-enriched cultures treated with CP1 (5 µM). (H) Regression analysis of MSN time courses shown in panels E-G, derived from long-read instability indices (CAG/day ± standard error; n = 3 independent biological replicates). Together, these data show dose-dependent slowing of somatic CAG repeat expansion by CP1 across dividing (U2OS) and post-mitotic (MSN-enriched) systems.

We next looked to assess the effect of pharmacological blockade of MutSβ DNA binding in a non-dividing system. We utilized the HD iPSCs carrying an expanded *HTT* CAG repeat (>125 CAG) described above (39). HD iPSCs were differentiated to striatal cultures enriched for medium spiny neurons (MSNs) (DARPP32+, FOXP1+, MAP2+), a neuronal subtype known to be vulnerable in HD (Figure S2). Cells were treated at maturity (day 36) with media containing vehicle (DMSO) or CP1 for 12 weeks, with media changes every 3-4 days. Minimal toxicity was observed with this compound at 5 µM over 12 weeks (Figure S2). CP1 treatment (5 µM) significantly slowed CAG repeat expansion in HD striatal neurons compared to vehicle (Figure 5).

## Discussion

Recent genetic studies have shown somatic expansion of the CAG repeat is the key pathogenic process driving HD onset and progression (5). Recognition of IDLs, lesions prone to form within the CAG repeat, is thought to be the primary event in the expansion process. In vitro experiments suggests this is mediated by MutSβ (MSH3/MSH2) and a wealth of evidence supports this MMR pathway as a major driver of SI and by extension HD pathogenesis (9,10,28,40). This makes MSH3 a protein of great interest both in terms of the mechanism of action at the CAG repeat and as a potential therapeutic target. To study MSH3 function we have developed a U2OS cell model in which MSH3 KO and complementation through a FlpIn site allow endogenous MSH3 function and the function of our introduced WT and mutant MSH3 transgenes to be compared. In addition, stable introduction of an HTT exon 118Q construct with a long CAG repeat (12) allows MSH3/CAG repeat interactions to be studied.

One important feature of this model is the ability to precisely regulate transgene expression through the TetOn system (12). This allows myc MSH3 6a (with MSH3 WT sequence) expression at a range of levels. At endogenous levels, expression of myc MSH3 6a restores MutSβ driven MMR activity and CAG repeat expansion (Figure 1). Protein and DNA interactions of the myc MSH3 6a are also similar to the endogenous protein (Figure 2 and 3). Interestingly, overexpression of the transgene led to an increase in MutSβ formation, MLH1 binding and CAG repeat expansion above that seen in WT cells, approaching the rate in FAN1 KO cells (Figure S1) (12). This shows overexpression of MSH3 results in an increase in active MMR complex formation that is capable of enhancing CAG repeat expansion, even in the presence of endogenous FAN1. This supports the idea that the expansion rate is determined by the balance between opposing factors, those that promote expansion (MSH3/MMR) and those that slow it down (FAN1), as suggested by recent in vitro work (41).

Having established a model in which transgenic MSH3 expression can recapitulate *MSH3* gene function we have looked in detail at the IDL binding pocket in MSH3 and its effect on CAG repeat expansion. Structural studies have identified tyrosine 245 and lysine 246 as key residues making close contact with DNA lesion (25). Substitutions at these positions (Y245S/K246E) in MSH3 abolish the interaction of MutSβ with IDLs, hairpin loops, and G4/R-loops(33,34) *in vitro*, suggesting this site could play a role in CAG repeat expansion. These corresponding substitutions were introduced into our myc MSH3 6a construct (as Y254S/K255E). myc MSH3^Y254S/K255E^ protein expression and MSH2 interactions were not obviously affected by the substitutions, but MLH1 binding tended to be reduced (Figure 2). MSH3^Y254S/K255E^ DNA binding was almost abolished, consistent with published *in vitro* data (33,34) (Figure 2). Thus, despite normal MutSβ formation, DNA binding was compromised. This may underlie the reduced MLH1 interaction as MutL recruitment to MutSβ would take normally place on a DNA substrate in situ.

Inefficient DNA interaction is also likely to underlie the MMR deficiency seen in myc MSH3^Y254S/K255E^ complemented cells. However, this did not reach the levels seen in KO or MSH3^E976A^ cells (Figure 1D). Importantly, CAG repeat expansion was impaired in MSH3^Y254S/K255E^ cells. The HTT exon 118Q CAG repeat was essentially stable over the 40 day time course. Similar cells carrying myc MSH3^E976A^ construct, a well characterised Walker B ATPase mutant, also failed to support CAG repeat expansion, and showed a tendency to small reductions in repeat length. We have seen this phenomenon previously in cells with compromised MSH3 function (39). These small differences between the E976A and Y254S/K255E forms in both EMAST and CAG repeat expansion assays may indicate some residual MMR activity is present in the DNA binding mutant.

To further study the role of the MSH3 IDL binding site in CAG repeat expansion we used a recently identified small molecule that binds close to Y254/K255. This molecule, referred to here as CP1, binds irreversibly to cysteine 252 within the MSH3 IDL binding pocket and inhibits DNA interaction (35). Using a CAG2 loopout oligo binding assay and ChIP we confirmed CP1 reduces MSH3 DNA binding *in vitro* and showed its inhibitory activity is retained in live cells (Figure 4). Remarkably, CP1 treatment of U2OS HTT exon 118Q cells slowed CAG repeat expansion in a dose dependent manner that reflected its inhibition of MSH3/DNA interaction (Figure 5).

While U2OS cells transduced with an exon1 construct have provided a useful tool to dissect the molecular mechanisms underlying CAG repeat expansion they may not reflect the full *in vivo* situation (ie somatic expansion in post mitotic neurons in the HD brain). Therefore, we have looked at the role of the MSH3 IDL binding pocket in our HD iPSC 125Q derived MSN enriched cultures, a physiologically relevant cell model. These predominantly neuronal post mitotic cultures (>95%) show increases in CAG repeat size over time in culture have been used previously to study the role of MMR proteins in SI (28,39). We treated differentiated cultures with the small molecule CP1 and assessed target engagement and measured repeat size over 12 weeks (Figure 5). In agreement with the effect observed in U2OS cells, CP1 treatment reduced MSH3 CAG2 loopout oligo binding and slowed expansion relative to control cells. No effect cell on morphology or viability was observed indicating CP1 is well tolerated by the cells at this concentration (Figure S2). Showing this effect in differentiated neuronal cells is important because it demonstrates CP1 is effective in slowing expansion of the *HTT* CAG repeat in non-dividing cells that express endogenous MSN DDR proteins.

The data reported here show MSH3 interactions with IDLs through the Y254/K255 binding pocket play an important part in the expansion process in U2OS HTT exon1 118Q model. We also report on a small molecule targeting this site that slows repeat expansion in U2OS HTT exon1 118Q cells and in HD patient stem cell derived neurons. CP1 is a covalent modifier that shows toxicity at higher concentrations and some promiscuity in terms of target engagement (35) and therefore may not represent a great therapeutic candidate in itself. However, our data suggests that the IDL binding pocket of MSH3 may represent a viable therapeutic target worth future study, particularly as *MSH3* is relatively tolerant of loss-of-function variation in humans (1).

## Acknowledgements

We would like to thank Brinda Prasad and Michael Finley, of the CHDI foundation, for critical review of the manuscript

## Author Contributions Statement

RG designed and performed experiments, interpreted results and wrote the paper with input from all authors; JD and FG designed and performed HD MSN experiments; ME prepared the biotinylated oligos and performed fragment analysis on U2OS samples; JH and LC performed fragment analysis on U2OS samples; PG performed long read sequencing; MF performed long read sequencing, designed software and performed statistical analysis of MSN samples; SJT supervised the project and obtained funding

## Data availability

The data are available in the article, in its online Supplementary data, or available upon request.

