## Supplemental figures S1 and S2 for "Genetic or pharmacological disruption of the MSH3 Y245/K246 IDL binding pocket slows CAG repeat expansion"

Supplementary data

Supplementary figures

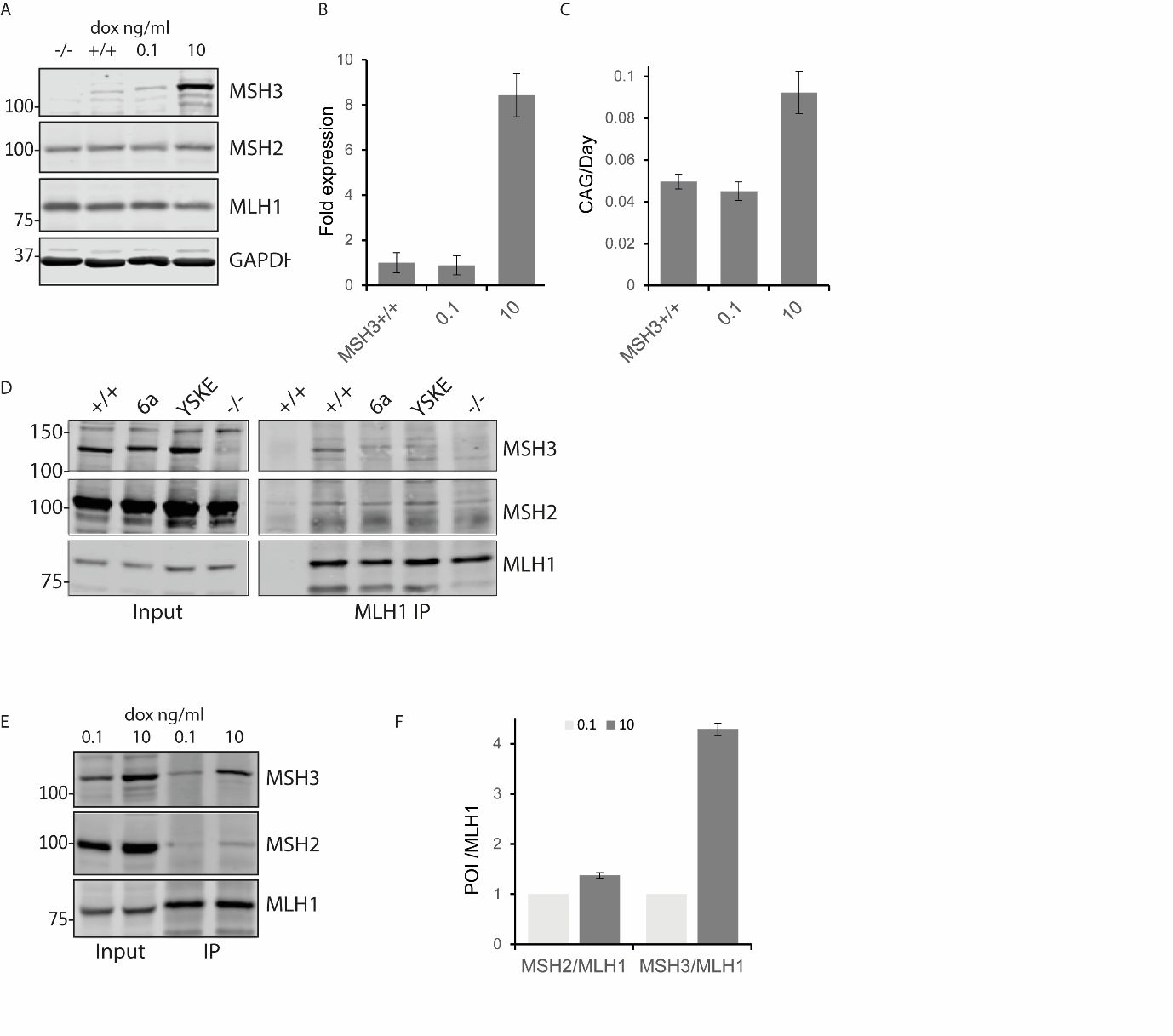

**Figure S1 MSH3 overexpression speeds up CAG repeat expansion**

(**A**) Immunoblots from U2OS WT (+/+), MSH3 KO (-/-) or KO cells complemented with myc MSH3 6a lysates from cells treated with vehicle or the indicated concentrations of dox. myc MSH3 6a expression close to endogenous levels is induced by 0.1 ng/ml dox, increasing dox concentration to 10 ng/ml leads to MSH3 overexpression (~8 fold) and MSH2 levels increase. (**B**) Quantification of blots normalized to WT MSH3 is shown in the histogram (n=2 independent experiments ± sd). (**C**) CAG repeat expansion rates from cells treated as in panel A. Note myc MSH3 overexpression leads to expansion rates above that seen in WT cells. (**D**) U2OS cell extracts from WT (+/+), MSH3 KO (-/-) or KO cells complemented with myc MSH3 6a or myc MSH3 YSKE and treated with 0.1 ng/ml dox were incubated O/N with anti-MLH1 antibodies and prot A/G magnetic beads. Beads were isolated and washed with a magnetic device. Input (5%) eluted proteins were imunnoblotted with the indicated antibodies. Low levels of MSH2 and MSH3 can be detected in the WT, myc MSH3 6a MLH1 IP fractions. The ratio of MSH3/MLH1 recovered in the IP fraction were similar from cells expressing endogenous MSH3 (WT) or myc MSH3 6a. MLH1 tended to pull down less myc MSH3 Y254S/K255E. (**E**) Immunoblots showing anti-MLH1 IP’s from U2OS extracts made from cells treated with 0.1 or 10 ng/ml dox concentrations, yielding the equivalent of endogenous or 8-fold myc MSH3 6a overexpression. Note myc MSH3 overexpression leads to an increase in MSH2 levels and a larger increase in MSH3/MSH2 (MutSβ) co-IP’d with MLH1. **(F)** Quantification of MSH2/MLH1 or MSH3/MLH1 in the IP fractions normalized to 0.1 ng/ml dox is shown in the histogram (n=2 ± sd).

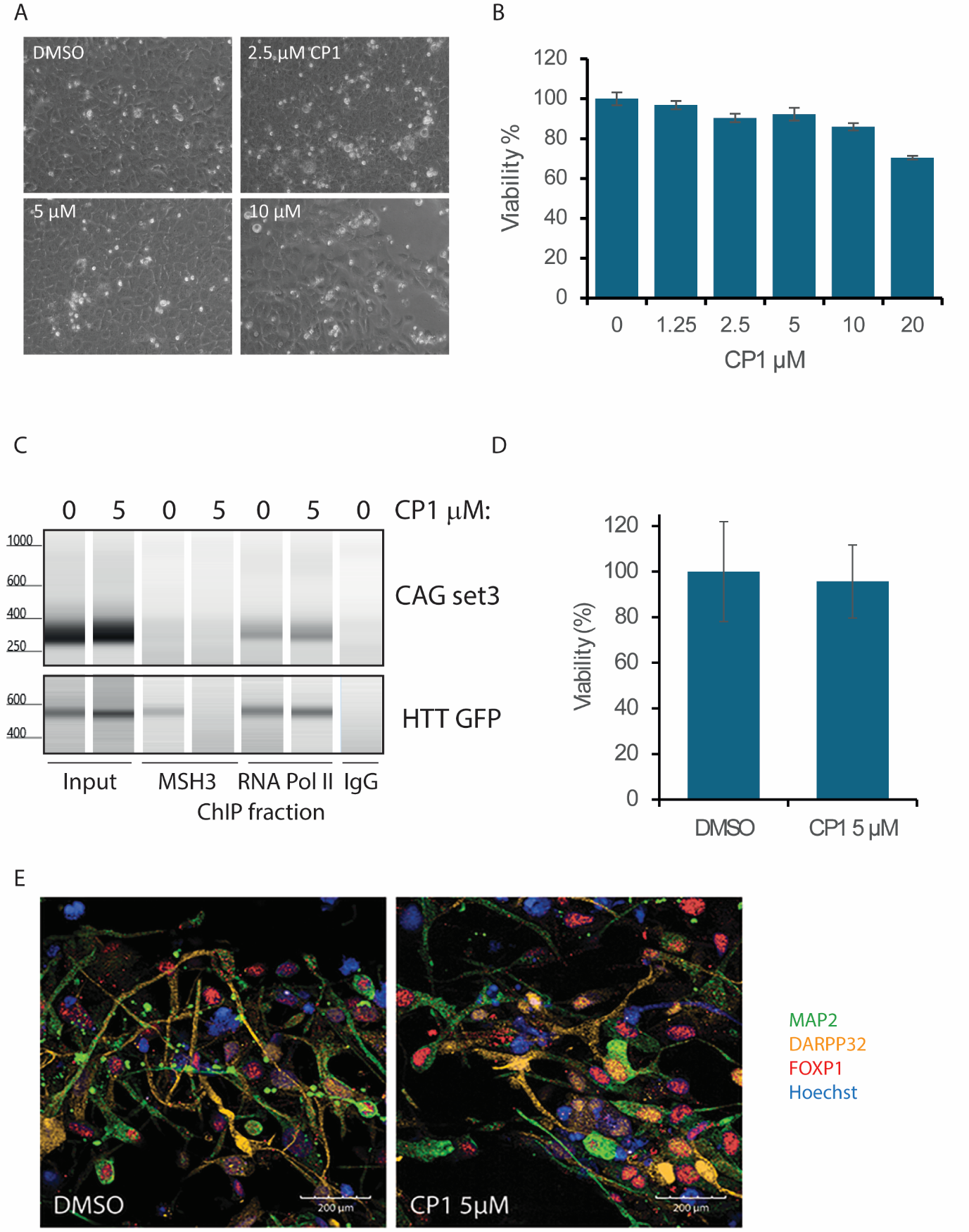

**Figure S2 CP1 toxicity to U2OS and iPSC 125Q derived MSN enriched cultures**

(**A**) Representative phase images of U2OS cells treated with the indicated concentrations of CP1. (**B**) MTT assay showing viability of U2OS cells following CP1 treatment. (**C**) FAN1 KO cells transduced with LV HTT exon 118Q were treated with CP1 5 µM (5) or vehicle (0). Cell extracts were prepared for ChIP analysis and immunoprecipitated with anti MSH3, RNA PolII or control non-specific mouse IgG antibodies. DNA from input and ChIP fractions was purified and probed with primers targeting the *HTT* exon 118Q CAG repeat present in LV HTT exon 118Q (CAG set 3) or GFP primers downstream from this in the insertion cassette (HTT GFP). Input (5%) and ChIP fractions were analysed using TapeStation apparatus and software. Although DNA levels in MSH3 ChIP fractions were low, CP1 treatment consistently reduced DNA levels in the MSH3 ChIP fractions (**D**) MTT assay showing viability in iPSC 125Q derived MSN enriched cultures following CP1 treatment (**E**) Immunocytochemistry of HD iPSC-derived striatal neurons after 12 weeks of vehicle (DMSO) or 5 µM CP1 treatment. Co-staining of cultured neurons for MAP2 (green), DARPP32 (orange) and FOXP1 (red).
